# Obtaining leaner deep neural networks for decoding brain functional connectome in a single shot

**DOI:** 10.1101/2020.04.22.056382

**Authors:** Sukrit Gupta, Yi Hao Chan, Jagath C. Rajapakse, the Alzheimer’s Disease Neuroimaging Initiative

## Abstract

Neuroscientific knowledge points to the presence of redundancy in the correlations of brain’s functional activity. These redundancies can be removed to mitigate the problem of overfitting when deep neural network (DNN) models are used to classify neuroimaging datasets. We propose an algorithm that removes insignificant nodes of DNNs in a layerwise manner and then adds a subset of correlated features in a single shot. When performing experiments with functional MRI datasets for classifying patients from healthy controls, we were able to obtain simpler and more generalizable DNNs. The obtained DNNs maintained a similar performance as the full network with only around 2% of the initial trainable parameters. Further, we used the trained network to identify salient brain regions and connections from functional connectome for multiple brain disorders. The identified biomarkers were found to closely correspond to previously known disease biomarkers. The proposed methods have cross-modal applications in obtaining leaner DNNs that seem to fit the data better. The corresponding code is available at https://github.com/SCSE-Biomedical-Computing-Group/LEAN_CLIP.

## 1. Introduction

Deep neural networks (DNN) have been successfully applied to a wide variety of classification problems, ranging from image recognition to language processing [21, 28]. In these applications, the choice of network architecture and number of trainable parameters affects network performance significantly. Larger networks learn more complex and accurate mappings, which comes with increased computational complexity and possibilities of overfitting [48]. On the other hand, having insufficient number of parameters limits the network’s ability to learn the correct mapping [60]. Therefore, finding the optimal neural network architecture is an important problem. One possible solution involves performing elimination on a large neural network by removing redundant nodes and connections. This has been shown to produce comparable performance with a smaller network but with better generalization capability [1, 4].

Neural networks and machine learning techniques have also been successfully applied to neuroimaging data. Non-invasive neuroimaging techniques are used to characterize functional and structural anomalies in the brain and aid in better diagnosis and treatment [13]. Functional and/or structural connectivity derived from these techniques are used as features for the neural network models to classify diseased and normal subjects [7, 15]. However, such studies typically involve many more input features (∼10, 000) than subject scans (∼1, 000), making the neural networks prone to overfitting. Also, experimental noise introduces systematic connectivity [34], which should be ignored during the classification task. Most crucially, overfitting is exacerbated by the presence of redundancies: not all functional connectivity features are important for differentiating between scan samples from two subject groups.

In recent works [16], the authors used feature salience scores to remove less salient features. In order to find salient features, DeepLIFT [44] was used to compute salience scores of both input features and hidden layer nodes. DeepLIFT is one of the recent attempts (besides Integrated Gradients [50], Layerwise Relevance Propagation [5] and SHAP [30]) that circumvent the issues with gradient-based approaches (such as zero gradients or discontinuities). These approaches find the contribution of nodes at each layer by propagating the contributions from the output layer. A major differentiating factor of DeepLIFT is that it gives consideration to both negative and positive contributions and computes the relevance scores efficiently in a single pass. By recursively removing the least salient features, these recent works arrived at a much smaller model with comparable accuracy to the original model, reducing the problem of overfitting without compromising on classification accuracy.

However, one limitation of these previous approaches is efficiency: they involve multiple elimination and fine-tuning steps to arrive at a smaller neural network. In this paper, we propose an algorithm called Layerwise Elimination of Accessory Nodes (LEAN) that performs the elimination of accessory nodes in a single shot. Accessory nodes are defined as nodes which do not influence the classification task significantly such nodes will be assigned a low importance score by a valid decoder. LEAN uses node salience scores derived from a valid DNN decoder to eliminate accessory nodes in the network, producing a leaner, more computationally efficient, and more generalizable deep neural network that is derived in a single shot.

LEAN leads to a drastic reduction in parameters, including the removal of a large number of input features. While powerful and efficient, this also leads to the loss of whole groups of correlated features. In neuroimaging datasets, such correlations exist due to the *specialization* property of the brain, whereby distinct sub-systems in the brain perform specialized tasks [17]. To tackle this, we introduce another algorithm called Correlation-based eLimination of InPuts (CLIP) that identifies and retains a subset of correlated input features. When these features are combined with the remaining input features from LEAN, it leads to an improvement in the DNN performance as compared to LEAN. Combining LEAN and CLIP leads to a model that provides the best of both worlds: low number of parameters with minimal accuracy drop.

Besides the goal of getting an efficient classifier with the highest accuracy and generalization ability, a relevant and important research direction in neuroscience is to find the biological markers that are associated with a brain state (disease or cognitive task), which differentiate it from another brain state. We denote identification of brain regions and connection that are associated with a particular brain state as brain decoding. Traditional methods for brain decoding include multivariate pattern analysis [18], sparse networks based feature selection [38], latent Dirichlet allocation [40], and the use of nodal features [43]. Such methods are however based on simple or linear models. On the other hand, DNN-based approaches build deep and hierarchical models and represent key patterns underlying brain activation in its nodes and connection weights [38]. Deep learning techniques make no assumptions about application specific a priori knowledge and therefore give consistent and unbiased decoding based on neuroimaging data. Recent applications of neural networks on neuroimaging data have derived the importance of input features in the classification task to uncover biomarkers for neurological diseases [16, 19] or reveal task-related brain functional modulations [27]. Identifying disease-specific biomarkers aids in building models and classifying and predicting the progression or occurrence of unknown diseases in individuals. To do so, we use the salience scores given by DeepLIFT to determine salient brain connections and regions for Alzheimer’s disease (AD), mild cognitive impairment (MCI), attention deficit hyperactivity disorder (ADHD), major depressive disorder (MDD), and Autism Spectrum Disorder (ASD) patients using their functional MRI scans.

In sum, we have made the following novel contributions in this work:

- We proposed an efficient layerwise node elimination approach, LEAN, that removes accessory input features and hidden layer nodes from a DNN in a single iteration. We further adapted it for brain decoding by proposing CLIP to add in a subset of correlated features, leading to a model with minimal number of connection while maintaining the accuracy.
- The proposed approach finds salient functional connections and identifies brain regions that are responsible of classification of brain disease.
- The proposed methods were applied on multiple brain disorders for classification of patients from healthy controls and identification of disease biomarkers.

## 2. Methods

### 2.1. Feedforward DNN

Given a pair (*x, d*) where *x* = (*x*_*i*_) denotes the input feature vector and *d* ∈ *D* denotes the sample label, we used a feedforward neural network of *L* layers with the first *L* −1 layers having rectified linear unit activation and a softmax layer at the end of the network. Let the weights and biases of the layer *l* be given by *W*_*l*_ and *b*_*l*_, respectively. The output of layer *l* ≠ {0, *L*} is given by:

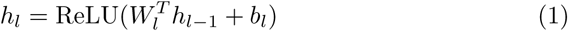

For the input layer, *h*_0_ = *x*. For the softmax layer, the output probabilities *y* of the input *x* belonging to class *k* is given by:

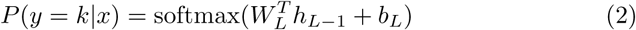

where *k* ∈ {1, …, *K*} represents the class label and the output layer weight *W*_*L*_ = [*w*_*k,L*_] and bias *b*_*L*_ = (*b*_*k,L*_). To learn the parameters of the network, the cross-entropy cost is defined as:

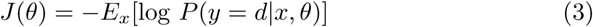

where *E*_*x*_ is the expectation taken over all scan samples *x* and 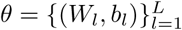 denotes all the parameters in the network.

### 2.2. Saliency of nodes in network layers

Let *f* be the neural network function mapping input *x* to output *y*. A simpler *explanation model g* is found such that *g* is both interpretable and an approximation of the model *f*. Let the number of nodes in layer *l* be *n*_*l*_. Using an appropriate reference, let us assign to each neuron *i* in layer *l* a contribution 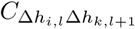 to the change in the output of neuron *k* in layer *l* + 1:

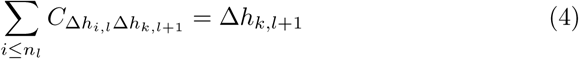

when *l* ∈ {0, …, *L* − 2} and

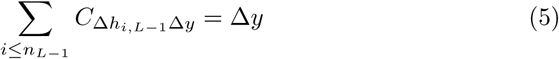

when *l* = *L* − 1, where Δ*h*_*i,l*_ is the change in the activation of the *i*^*th*^ neuron of layer *l* due to the input relative to the reference. 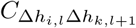 is computed from the Linear, Rescale and RevealCancel rule from [44].

Given the reference input 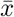 and the original input *x*, we substitute Δ*y* = 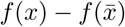 and *g*(*x*) = *f* (*x*), giving us an equation for the model *g*:

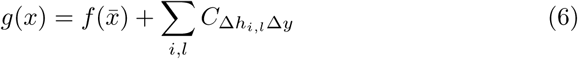

where the contribution of nodes in each layer *l* to the output *y* is given by 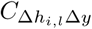 [44]:

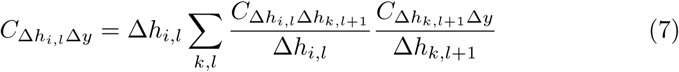

Contribution of nodes in all layers 0 ≤*l* < *L* to the output *y* is obtained by backpropagating contributions of layers to the output. For nodes in each layer *l* ∈ {0, …, *L* − 1}, we compute the *salience score vector c*_*l*_ given by:

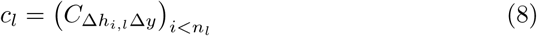

where 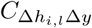 is derived from Equation 7. The DeepLIFT method [44] computes the layer’s salience scores based on change in the output from a reference, allowing information to propagate across the network layers even when the gradient is zero. We compute the average contributions for all layers over multiple test samples to get the final salience score for each feature.

### 2.3. LEAN: Layerwise elimination of accessory nodes

A decoder can identify a subset of distinguishing input features and weights that are important to classify samples. We proposed a brain decoding strategy in [16], where we obtained the salience scores for input features and hidden layer nodes. In the proposed scheme, a fraction *µ* of nodes with the lowest salience scores *c*_*k*_ from layers *k* ∈{0, …, *L* −1} are removed iteratively and the pruned model is fine-tuned at each step. Although the proposed strategy improved the classification performance, the process involved multiple iterations of elimination and fine-tuning that is time-consuming and cumbersome. Thus, we propose an efficient strategy to find a layerwise salience score threshold, below which nodes from the layers (input and hidden) of the DNN are removed.

For the salience score vector *c*_*l*_ in each layer, we determined the best fit distribution by computing the log likelihood of the model fit. We did not make any assumption about the best fit distribution and used multiple distributions (viz; power-law, log-normal, exponential and stretched exponential) to find the best fit for the salience score distribution at each layer. The power-law distribution is defined by the distribution of the salience scores *c*_*il*_ for nodes *i* < *n*_*l*_ in layer *l* [2, 10]:

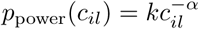

where *α* is the scaling parameter, 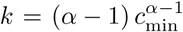, and *c*_min_ is the minimum degree that obeys the power-law. The distribution parameters were computed using maximum likelihood estimation [10]. We compute statistical significance for the salience scores of nodes in their respective layers and remove the nodes with *p* −*value* ≥ 0.95. The complete overview of the layerwise elimination of nodes is given in Algorithm 1.

#### Algorithm 1 LEAN: Layerwise elimination of accessory nodes

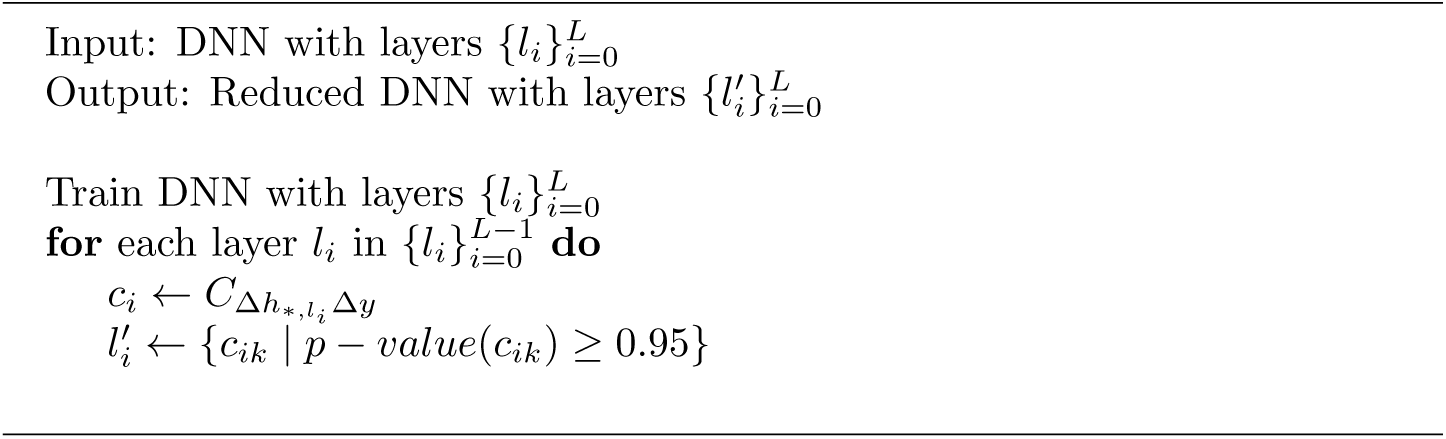

### 2.4. CLIP: Correlation-based elimination of inputs

Besides the presence of a large number of correlated features, brain functional connectomes are also known to be modular in nature [17, 47] and such modularity or clusters give rise to correlated features. Different modules correspond to different sub-systems in the brain, performing a specific function. Examples of these modules are shown in Figure 1(a). Modules or clusters have stronger connectivity between nodes within the module and weaker connectivity with nodes outside the module. This gives rise to the intuition that the functional connectivity for some regions of interest (ROI) - especially ROIs within the same module - are correlated with each other.

**Figure 1:**
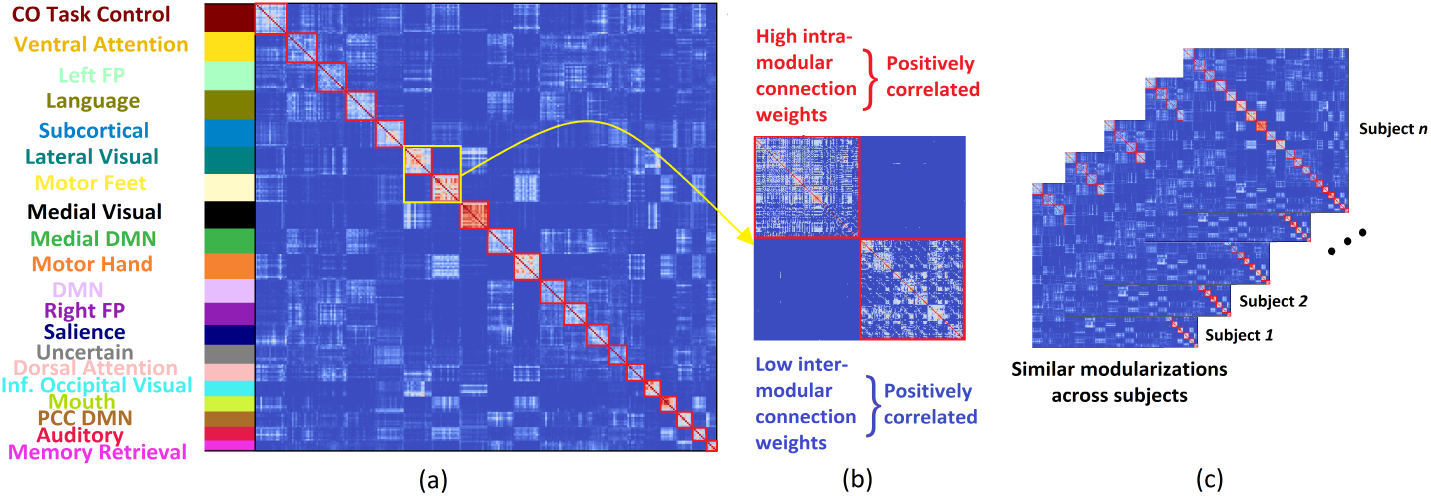
Visualization of the functional connectivity matrices. (a) The matrix has its rows rearranged such that nodes in the same functional clusters are placed consecutively. The labels of different functional modules are given by the left color bar. Image from [17]. (b) A magnified view of a subset of the correlation matrix is provided. Cells colored white have a higher weight than the cells colored blue. The white patches along the left diagonal represent intra-cluster connections, while the blue patches along the right diagonal represents inter-cluster connections. Intra-cluster connections have high and inter-cluster connections have low values across subjects and are therefore positively correlated. (c) A schematic of how some of the clusters are similar across subjects.

Although some modules involved in cognition and decision-making vary across subjects, others involved in functions related to perception and motor control are stable across subjects [17, 32]. This gives rise to inter-subject relationships among functional connectivity features. While the extent of how high correlations is difficult to determine, the correlation between pairs of inter-cluster connections would likely be lower than the correlation between pairs of intra-cluster connections. This is because of the sparser and variable inter-cluster connections, and the relatively denser and more stable intra-cluster connections due to similar modularizations as shown in Figure 1(b) and (c). For example, the dense connections within the visual module [17, 32], which has a low inter-subject variability, should be highly correlated across subjects.

#### 2.4.1. Finding clusters of correlated features

While there are other dimensionality reduction approaches such as Principal Component Analysis or Independent Component Analysis that reduce the problem of multicollinearity [22], such approaches require either prior knowledge of the number of components to be used, or extensive experimentation to arrive at the optimal number. Alternatively, a clustering-based approach presented below provides a more principled solution. Figure 2 provides a schematic of the intuition behind finding such clusters that are spread across the distribution of salience scores.

**Figure 2:**
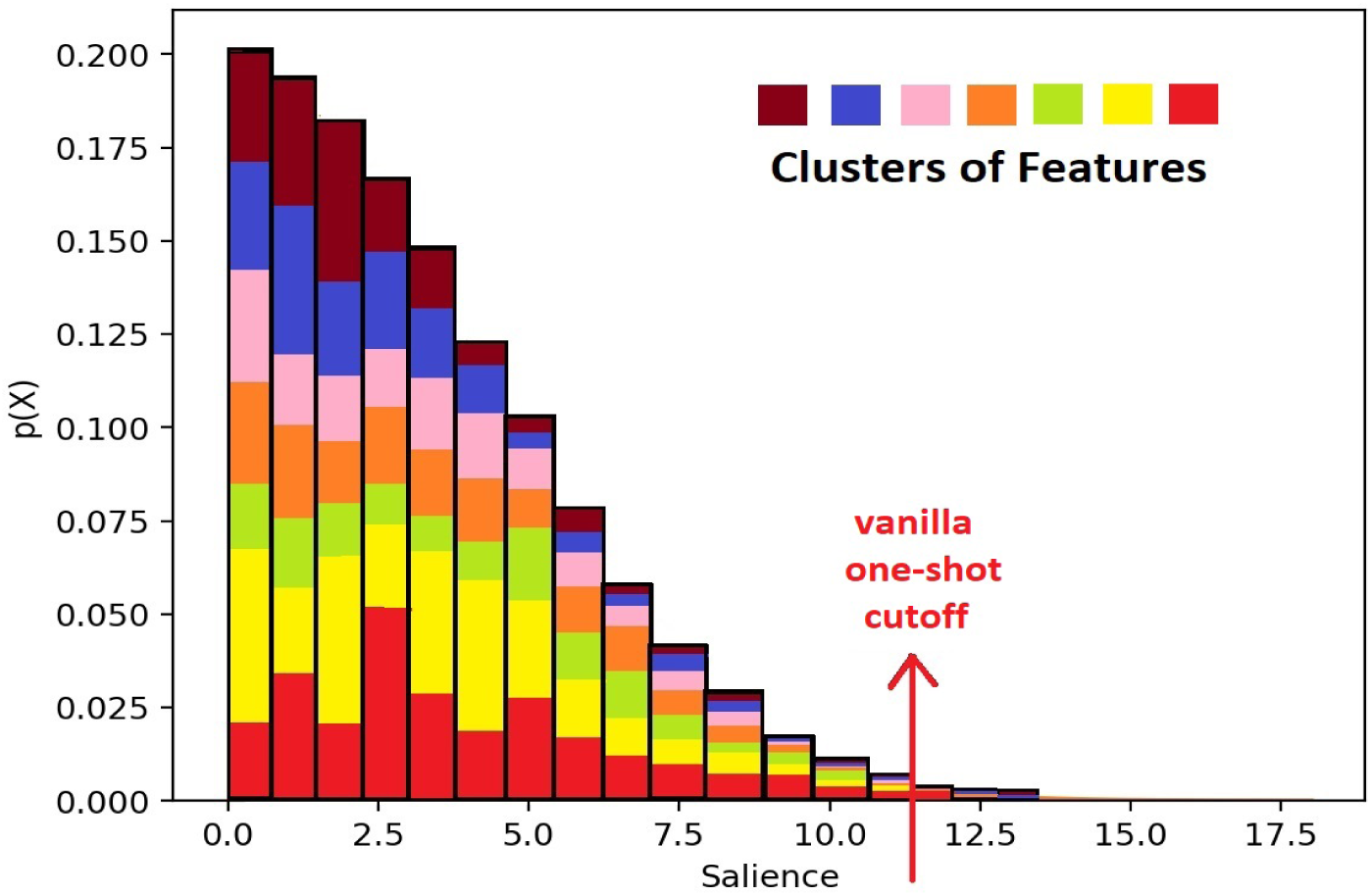
Schematic of how the features are distributed in different clusters throughout the distribution of salience scores and how salience scores consider only features from a few clusters.

We formulate this problem in terms of finding clusters/communities of correlated features, where we compute the similarity matrix, *S* = {*s*_*ij*_}, where *s*_*ij*_ denotes the correlation between features *i* and *j* from the training data:

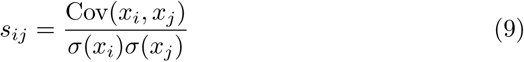

where Cov finds the covariance between two variables and *σ* finds the standard deviation, and *x*_*k*_ denotes the vector containing values of feature *k* across the samples.

We perform clustering of the similarity matrix *S* to obtain a cluster vector *m* = (*m*_*k*_) where each element *m*_*k*_ corresponds to the cluster label assigned to each voxel *k* from a set {1, 2, …, *M*} and *M* of clusters. This is done by the minimization of the normalized cut cost given by:

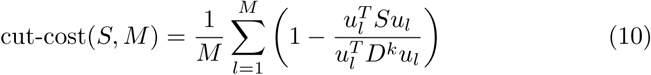

where *D* denotes the diagonal degree matrix of *S* and 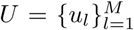 is a set of binary matrices representing *m*, such that *l*th module is given by 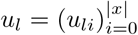 where *u*_*li*_ = 1(*m*_*i*_ = *l*) and 1(·) denotes the identity function.

The minimization of the cut-cost in (10) is performed by multi-class spectral clustering [49]. The number of clusters is found by computing the elbow point [42] (i.e. the point of maximum curvature) of the scree plot of the eigenvalues of similarity matrix *S*. However, since the number of features |*x*| are often large, the similarity matrix has a large size and computing its eigendecomposition becomes expensive. Therefore, we sparsified the matrix *S* by thresholding correlations with values less than 0.3 (which is a recognised threshold for low correlation values [20]). Thereafter, we used the Implicitly Restarted Arnoldi Method [29] to perform eigendecomposition on the sparse matrix to obtain the eigenvalues.

#### 2.4.2. Selecting subsets of correlated features

For each of the computed clusters, a subset of features that have the highest correlation with features from the same cluster was selected. We do this by using the *intra-cluster degree, α*_*i*_, of feature *i* given by

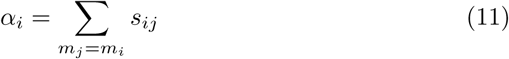

We select a fraction of nodes with the highest *α*_*i*_ values from each cluster and include them as features in addition to the ones selected from one-shot elimination pruning. The whole process for removal of subsets of correlated features is summarized in Figure 3 and Algorithm 2.

**Figure 3:**
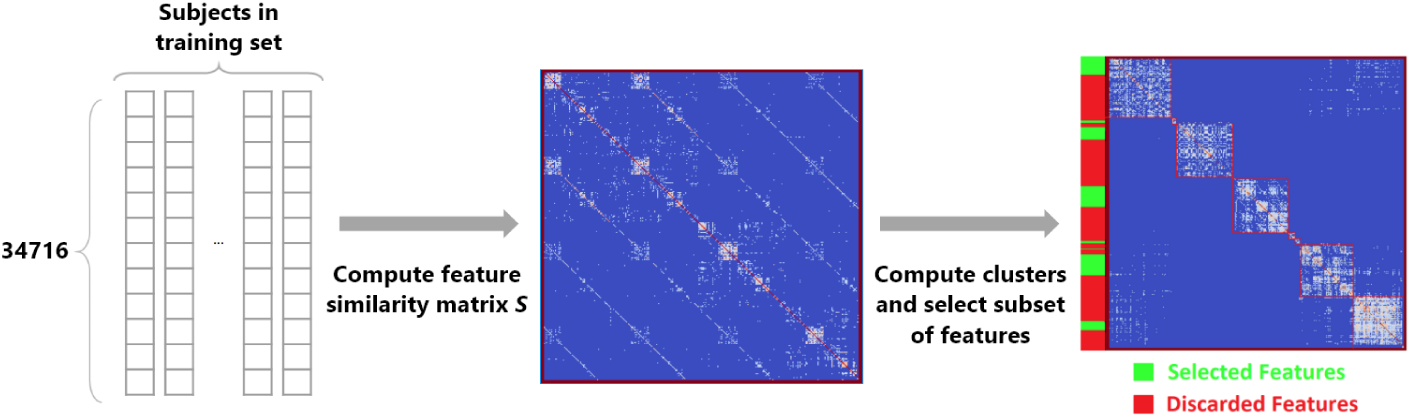
Overview of the feature selection process. To generate *S*, the vector of features representing each subject is concatenated and the similarity matrix is computed. Thereafter, clustering is performed on *S* and a subset of nodes with the highest intra-cluster degree each cluster are retained, while the rest are discarded. The matrix on the right has been rearranged such that nodes in the same cluster are placed consecutively.

### 2.5. Combining LEAN and CLIP

We argue that the presence of correlated features leads to a sudden drop in accuracy with LEAN because while performing recursive elimination, we retain a fraction of features from clusters of correlated features at different thresholds. However, with LEAN, entire clusters of correlated features are lost at once. This problem has been studied previously for various linear and non-linear classifiers [62] by using different feature selection methods [54]. For instance, in ordinary least square regression, the existence of multicollinearity leads to a larger standard error [14] and affects the interpretation of feature salience - a feature that would have been deemed as important is no longer significant when another correlated feature is present. We hypothesize that LEAN discards too many input features at once and using the additional subsets of correlated features (identified by Algorithm 2) will improve the model accuracy. Thus, while hidden layers nodes are eliminated only using LEAN (Algorithm 1), the input features are first eliminated using LEAN and then some are added back from the subsets of correlated features via CLIP (Algorithm 2).

#### Algorithm 2 CLIP: Correlation-based elimination of inputs

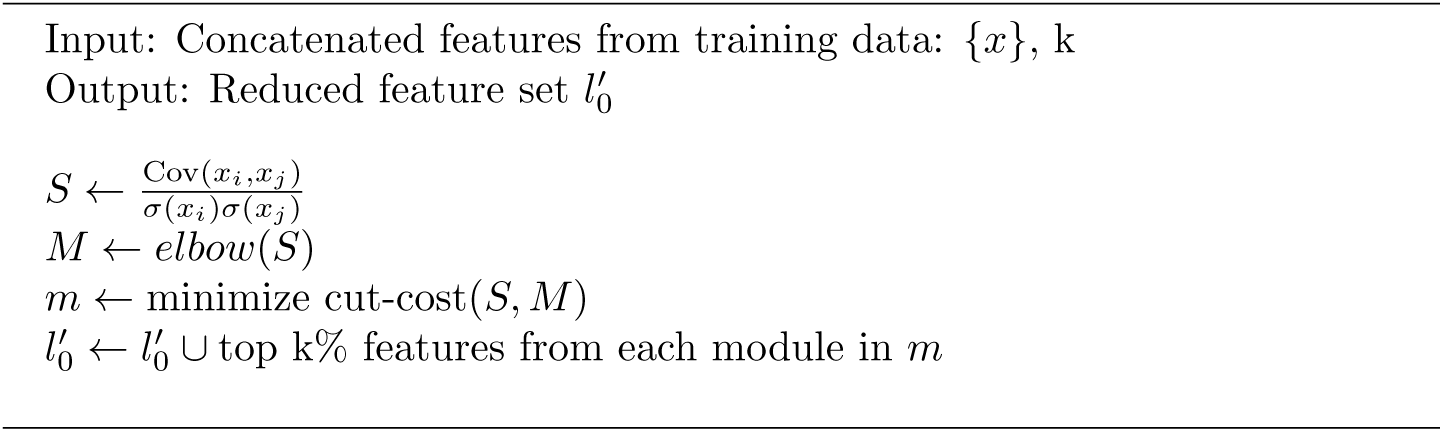

### 2.6. Decoding the brain functional connectome associated with brain disease

Brain decoding is defined as the identification of brain activity patterns that emerge from activations as well as interactions among specific brain regions and connections, which distinguish one brain state from another. Decoding the DNN trained on connectome features translates to identifying salient features of the DNN, which correspond to biomarkers (i.e. key brain connections and ROI) associated with the brain state. We demonstrated this approach in [16] for the first time by using feedforward DNN. We successfully adopted DeepLIFT in decoding brain functional connectivity in [16], which efficiently computes salience scores for input features in a single pass and then recursively eliminate irrelevant. Such an approach only focuses on input features and does not optimize the DNN architecture. In this paper, we improve upon our previous work by proposing a combination of LEAN and CLIP for not only to decode the input feature but also finds leaner DNN model for efficient classification without the loss of accuracy. By using LEAN and CLIP on resting-state fMRI brain scans gathered in brain disease, we achieve state-of-art accuracies for disease classification with leaner DNN models and salient input features. The decoded input features correspond to brain connections that are associated with brain disease.

## 3. Results

### 3.1. Datasets

We downloaded resting state functional MRI scans for AD and MCI from the Alzheimer’s Disease Neuroimaging Initiative (ADNI); for ADHD from the International Neuroimaging Datasharing Initiative (INDI) [6]; for MDD from the data provided by the Creativity and Affective Neuroscience Lab in the Brain Imaging Center of Southwest University and for ASD from the Autism Brain Imaging Data Exchange (ABIDE). The details of the acquisition protocols, subjects, preprocessing pipelines are attached in the supplementary materials.

### 3.2. Features for classification

We used the Power atlas [35] to obtain functionally diverse 264 ROIs spanning the entire cerebral cortex. Average time series were computed for voxels within a spherical radius of 2.5mm surrounding each ROI, and the functional connectivity matrix for each scan was generated by computing the Pearson correlation coefficient (without thresholding) for each pair of ROI. Since the matrices are symmetrical, we consider only the upper triangular connectivity matrix and flatten it into an input vector for the network. This resulted in an input vector with 34716 elements.

### 3.3. Classification for full feature set

Both the encoder and decoder were implemented in Python using the Keras, Tensorflow and DeepLIFT libraries. For datasets with more than one scan per subject, we ensured that all the subject scans were either in the training or test set. For all datasets, a batch size of 8 was used along with a learning rate of 10^−4^. The dataset was separated into train and test set at a 80:20 split.

Besides the feedforward neural networks (FFN), different convolution neural network (CNN) architectures [8, 31] and support vector machines (SVM) were implemented with different parameters. For CNNs, we gave the connectivity matrix as an input, and varied the number of filters and the number of layers. The number of filters in each layer was varied based on the number of weights in the corresponding FFN such that the number of trainable parameters were same in both the architectures. The number of trainable weights were changed from 1.7 × 10^4^ to 3.5 × 10^7^. For SVMs, we varied the parameters (*C* ∈{0.001, 0.01, 0.1, 1, 10}, *γ* ∈{0.001, 0.01, 0.1, 1}) and tried different kernels such as linear, polynomial, RBF and sigmoid. For the FFN, the number of hidden layers and the number of neurons in each hidden layer were varied. The final architecture for the full feature set was obtained using grid search from accuracies obtained from stratified 5-fold cross validation on the train set. We added dropout of 0.1 to the hidden layers and imposed early stopping to prevent overfitting. The details of the different configurations for the DNNs can be found in the supplementary materials. We observed that the highest accuracies were still given by FFN (except in case of CN vs AD, where the difference was insignificant). The average accuracies along with the standard deviation for the SVM, CNN and FFN models are reported in table 1 and the corresponding architectures are reported in table S5 in the supplementary materials. These accuracies are obtained from repeating the experiments using 10 different seeds. For each seed, 5-fold cross validation was performed.

**Table 1:** Performances of the best models of different models on the neuroimaging datasets.

| Disorder | SVM | CNN | FFN |
| --- | --- | --- | --- |
| ASD | 68.0% ± 3.3% | 70.6% ± 2.9% | 70.9% ± 4.2% |
| AD | 71.3% ± 6.5% | 78.1% ± 5.3% | 77.4% ± 4.6% |
| ADHD | 54.8% ± 5.5% | 60.5% ± 4.5% | 61.1% ± 4.2% |
| MCI | 64.6% ± 4.3% | 66.5% ± 3.8% | 67.5% ± 4.1% |
| MDD | 73.3% ± 3.9% | 77.1% ± 3.6% | 78.3% ± 3.2% |

### 3.4. Classification with feature subsets from CLIP and LEAN

We tried 3 different elimination approaches: ‘LEAN’, ‘CLIP + LEAN (Inputs Only)’ and ‘LEAN + CLIP’. ‘LEAN’ (described in Algorithm 1) involves keeping only statistically significant features at each layer; ‘CLIP + LEAN (Inputs Only)’ includes subsets of correlated features (Algorithm 2) in addition to DeepLIFT features, but no elimination is performed for the hidden layers; and ‘CLIP + LEAN’ goes even further to prune the hidden layers, as described in Section 2.5. For CLIP, we generated the similarity matrix (described in Section 2.4.1) to derive a subset of correlated features that are combined with features obtained from LEAN. We did this for each of the seeds and folds and retained 5% of the top features for each cluster. Table 2 summarises the changes in classification accuracy and remaining number of trainable parameters (compared to the the original model) for different approaches on the 3 datasets.

**Table 2:** Comparison of functional connectivity based classifiers. Params represent the number of parameters left in the reduced model, relative to the model that uses the full feature set.

|  | All<br>Features | LEAN |  | CLIP + LEAN<br>(Inputs Only) |  | CLIP + LEAN |  |
| --- | --- | --- | --- | --- | --- | --- | --- |
| Disorder | Accuracy | Acc | Params | Acc | Params | Acc | Params |
| ASD | 70.9% | 68.1% | 0.5% | 69.3% | 9.5% | 68.9% | 1.1% |
| AD | 77.4% | 75.8% | 0.7% | 78.1% | 7.7% | **78.1%** | 2.0% |
| ADHD | 61.1% | 59.0% | 2.1% | 59.3% | 9.9% | **59.4%** | 5.7% |
| MCI | 67.5% | 64.5% | 0.6% | 65.3% | 6.9% | **66.2%** | 2.0% |
| MDD | 78.3% | 75.2% | 1.2% | **76.4%** | 10.9% | 76.0% | 2.6% |

As seen in table 2, we observe that relative to the full feature set, LEAN is able to reduce the model parameters drastically to around 0.5% to 2.1% of the original number of parameters. Intuitively, keeping the most important features should lead to a model with minimal drop in accuracy. However, we found that the drop in accuracy is large (1.6% in ADHD to 3.1% in MDD) and thus LEAN cannot be directly used. One can argue that a different p-value threshold can be used to allow more features to be included, but it will take several iterations to arrive at the threshold which gives the optimal model. Furthermore, such a tuned threshold is unlikely to generalize to other situations.

We reduced the drop in accuracy by identifying subsets of correlated features in ‘CLIP + LEAN (Inputs Only)’. Evidently, our approach leads to a smaller drop in accuracy as compared to the LEAN approach (and even lead to an increase in accuracy for CN/AD). Although we obtain similar classification performance when we only prune the inputs (and not the hidden layers), the number of remaining parameters (7% to 11% of the original) are much larger than LEAN. By extending the node elimination process to the hidden layers in ‘CLIP + LEAN’, we further reduced the number of parameters involved to levels quite close to the LEAN approach, with similar accuracies (and with consistently higher accuracies than LEAN).

Comparing the results of ‘CLIP + LEAN (Inputs Only)’ and ‘CLIP + LEAN’, it is seen that there is no significant trend in terms of change in accuracy. This shows that we can safely remove hidden nodes to obtain a leaner model. Thus, our subsequent analysis will be focused on LEAN and ‘CLIP + LEAN’.

#### 3.4.1. Reduction in overfitting

Overfitting happens during model training when the test loss increases after reaching a minimum [60]. If a model overfits, it fails to generalise and remembers only the training data. To evaluate overfitting, we use the difference between the test set loss and train set loss at the epoch as evaluated by cross-entropy when the best model was found. Because of the small size of the datasets, the difference of train and test losses renders a stable measure of overfitting [33].

The model was first trained with early stopping, such that the best model was chosen by looking at the test accuracies of each of the epochs. The best model is chosen at the epoch where test accuracy is highest. Table 3 shows the extent of overfitting for each of the datasets with cross-entropy being the metric used to compute the loss. The reported scores were calculated by averaging over 5 folds. As shown in the table, there is a clear reduction of overfitting across all datasets for both LEAN and ‘CLIP + LEAN’, after these algorithms were used to generate the leaner model.

**Table 3:** The differences of test and train losses computed using cross-entropy.

| Disorder | All Features | LEAN | CLIP + LEAN |
| --- | --- | --- | --- |
| ASD | 0.50 | 0.28 | 0.20 |
| AD | 0.34 | 0.14 | 0.13 |
| ADHD | 0.29 | 0.06 | 0.01 |
| MCI | 0.13 | 0.002 | 0.02 |
| MDD | 0.61 | 0.31 | 0.36 |

#### 3.4.2. Comparison with other redundancy removal methods

We compared performances of ‘CLIP + LEAN’ with those of SVM and logistic regression, including their own feature importance methods. Experiments were conducted, keeping 1%, 5% and 10% of features as this is the same range of remaining features obtained from our proposed algorithms. For SVM, only the linear kernel is able to provide importance scores and for this comparison, the best model with a linear kernel was chosen for each dataset. As seen in table 4, as the number of features remaining increases, model accuracy for logistic regression generally increases but such a trend is less pronounced for SVM. Ultimately, ‘CLIP + LEAN’ still generally outperforms both the SVM and logistic regression models.

**Table 4:** Accuracies of models produced by keeping X% most important features from SVM and logistic regression models. C + L = CLIP + LEAN.

| Disorder | FFN | C + L |  | 1% | 5% | 10% | 100% |
| --- | --- | --- | --- | --- | --- | --- | --- |
| ASD | 70.9% | 68.9% | SVM | 53.0% | 59.0% | 67.7% | 68.0% |
|  |  |  | Logreg | 63.1% | 67.0% | 66.6% | 67.9% |
| AD | 77.4% | 78.1% | SVM | 66.8% | 70.2% | 71.6% | 71.3% |
|  |  |  | Logreg | 70.9% | 72.7% | 72.2% | 71.3% |
| ADHD | 61.1% | 59.4% | SVM | 53.2% | 51.6% | 52.0% | 53.1% |
|  |  |  | Logreg | 52.5% | 51.9% | 52.1% | 53.2% |
| MCI | 67.5% | 66.2% | SVM | 64.6% | 62.9% | 60.1% | 58.7% |
|  |  |  | Logreg | 56.1% | 57.9% | 58.4% | 58.7% |
| MDD | 78.3% | 76.0% | SVM | 69.4% | 72.5% | 72.9% | 73.3% |
|  |  |  | Logreg | 70.5% | 72.6% | 72.9% | 73.3% |

Additionally, since traditional models like SVM and logistic regression assume that the used features are uncorrelated, we used CLIP to retain a subset of uncorrelated features and compared the performance of the resulting models. As seen in table 5, the classification accuracy usually suffers when a feature subset is used. However, the accuracy is significantly lower (*p* −*value* < 10^−3^) than the corresponding FFN in all the cases (except for MCI).

**Table 5:**
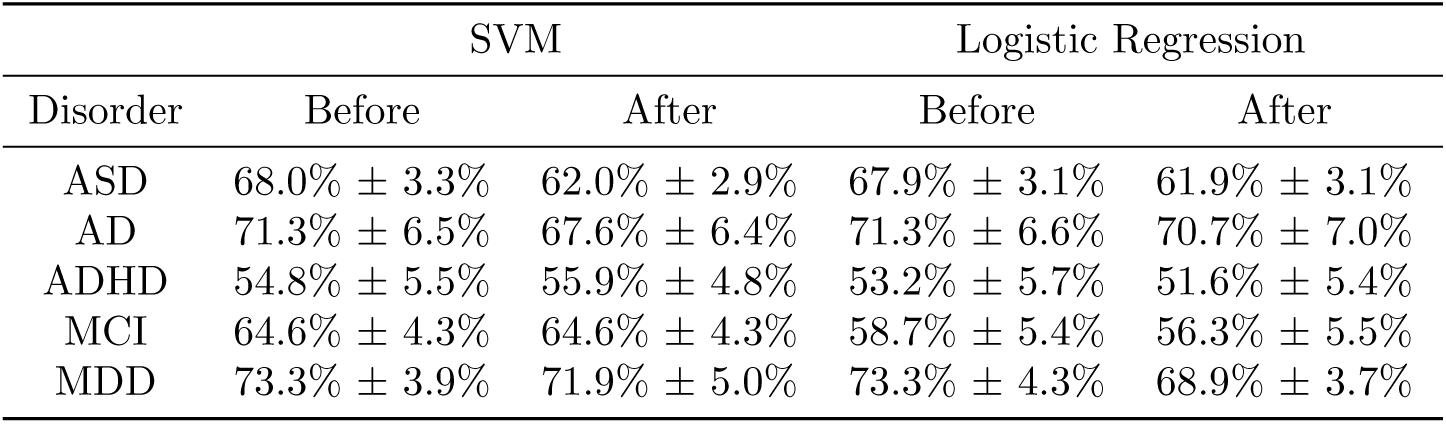
Accuracies of baseline models before and after CLIP was applied.

| Disorder | SVM |  | Logistic Regression |  |
| --- | --- | --- | --- | --- |
|  | Before | After | Before | After |
| ASD | 68.0% ± 3.3% | 62.0% ± 2.9% | 67.9% ± 3.1% | 61.9% ± 3.1% |
| AD | 71.3% ± 6.5% | 67.6% ± 6.4% | 71.3% ± 6.6% | 70.7% ± 7.0% |
| ADHD | 54.8% ± 5.5% | 55.9% ± 4.8% | 53.2% ± 5.7% | 51.6% ± 5.4% |
| MCI | 64.6% ± 4.3% | 64.6% ± 4.3% | 58.7% ± 5.4% | 56.3% ± 5.5% |
| MDD | 73.3% ± 3.9% | 71.9% ± 5.0% | 73.3% ± 4.3% | 68.9% ± 3.7% |

#### 3.4.3. Comparison of algorithm runtime

In order to evaluate time efficiency of different algorithms, average time taken per epoch to train the neural network were computed. As shown in table 6, all variants of our proposed redundancy reduction algorithms lead to a reduction of time taken - the increase being greater especially if the original ‘All Features’ model used was large like in the case of MDD. Here, we also specified the time taken for the one time operation to compute the feature clusters in CLIP. On average (over datasets and seeds), pre-processing took 135 minutes where the eigenvalue computation of the similarity matrix *S* is the source of the bottleneck. This can be significantly reduced by lowering the number of eigenvalues to be computed - in our experiments, we defined it to be 1000. Also, importance score computation took, on average, a range of approximately 20s for AD to approximately 2 minutes for MDD (which starts off from a large model). All experiments were performed using 4 x NVIDIA Tesla P100 16GB on a Linux server with 72 cores. Comparing the proposed algorithms with each other, there are slight variations in the timings but the differences are largely insignificant (except for the case of MDD) and both variants take approximately the same amount of time to run. We also concluded that the pruned models, LEAN and ‘CLIP + LEAN’, take a significantly shorter amount of time to run as compared to the full model (*p* −*value* < 10^−4^). When a relatively small model is used, both variants take approximately the same amount of time to run.

**Table 6:**
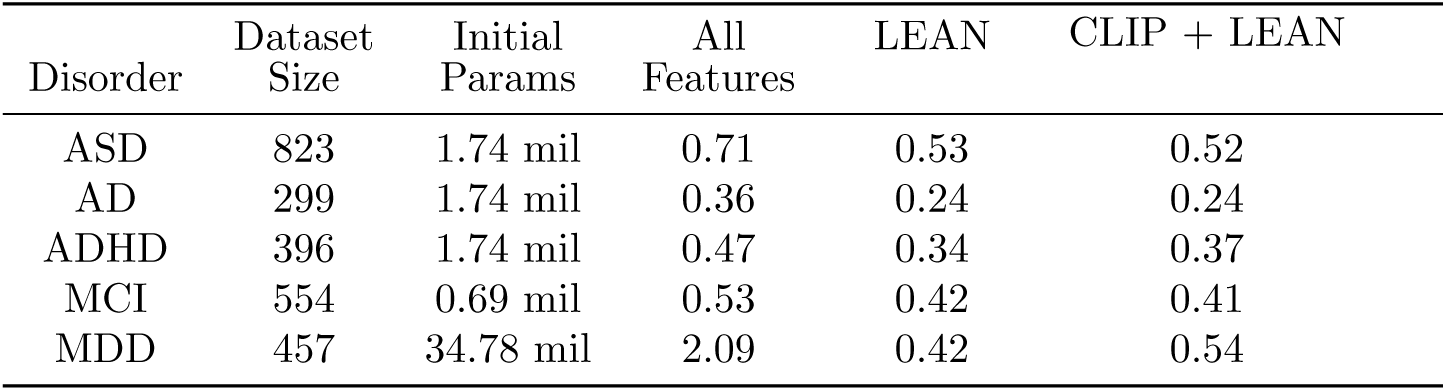
Average time taken per epoch (in seconds) for each variant of the proposed algorithms. mil = million.

| Disorder | Dataset Size | Initial Params | All Features | LEAN | CLIP + LEAN |
| --- | --- | --- | --- | --- | --- |
| ASD | 823 | 1.74 mil | 0.71 | 0.53 | 0.52 |
| AD | 299 | 1.74 mil | 0.36 | 0.24 | 0.24 |
| ADHD | 396 | 1.74 mil | 0.47 | 0.34 | 0.37 |
| MCI | 554 | 0.69 mil | 0.53 | 0.42 | 0.41 |
| MDD | 457 | 34.78 mil | 2.09 | 0.42 | 0.54 |

### 3.5. Salient brain features

Thus far, we have seen how salience scores help to reduce the number of parameters involved, leading to a leaner model with minimal reduction in accuracy (or at times, even leading to an increase in model accuracy). We also use the salience scores to identify important features and ROI for the different diseases. The input fed into our neural network model is a 34716-dimensional vector and each element in the vector represents a correlation score between a pair of ROI (i.e. strength of a functional connection). We analyse the importance of the functional connections by using the salience scores and the importance of ROI by computing the sum of salience scores of the functional connections incident on the ROI. Since the Power atlas [35] does not provide anatomical labels, the Crossley atlas [12] (which has anatomical labels) was used to map ROI from the Power atlas on the basis of Euclidean distance. Then, ROI with the top 10% highest scores are visualised using Nilearn.

From Figure 4b, we found that the salient connections for MDD lie between the transverse temporal gyrus and lingual gyrus; the transverse temporal gyrus and thalamus; superior occipital gyrus and precuneus, and between extra-nuclear and inferior temporal gyrus. For AD, we found that the connections between the superior temporal and subcallosal; superior temporal and extra nuclear, and between the uncus and inferior temporal regions were salient (Figure 4a). For MCI, however, we found that only the connections between the uncus, subcallosal and uncus inferior temporal were salient (supplementary figure S2(b)). For ADHD, the salient connections were found between the paracentral and superior temporal; paracentral and inferior frontal; inferior temporal and superior occipital; supramarginal and sub-gyral; and supramarginal and extra nuclear regions (supplementary figure S2(a)). For ASD, salient connections were found between regions in the frontal inferior orbital to the putamen, frontal superior orbital, and frontal inferior operculum; the putamen and temporal pole superior; and cerebellum and the vermis (supplementary figure S2(c)).

**Figure 4:**
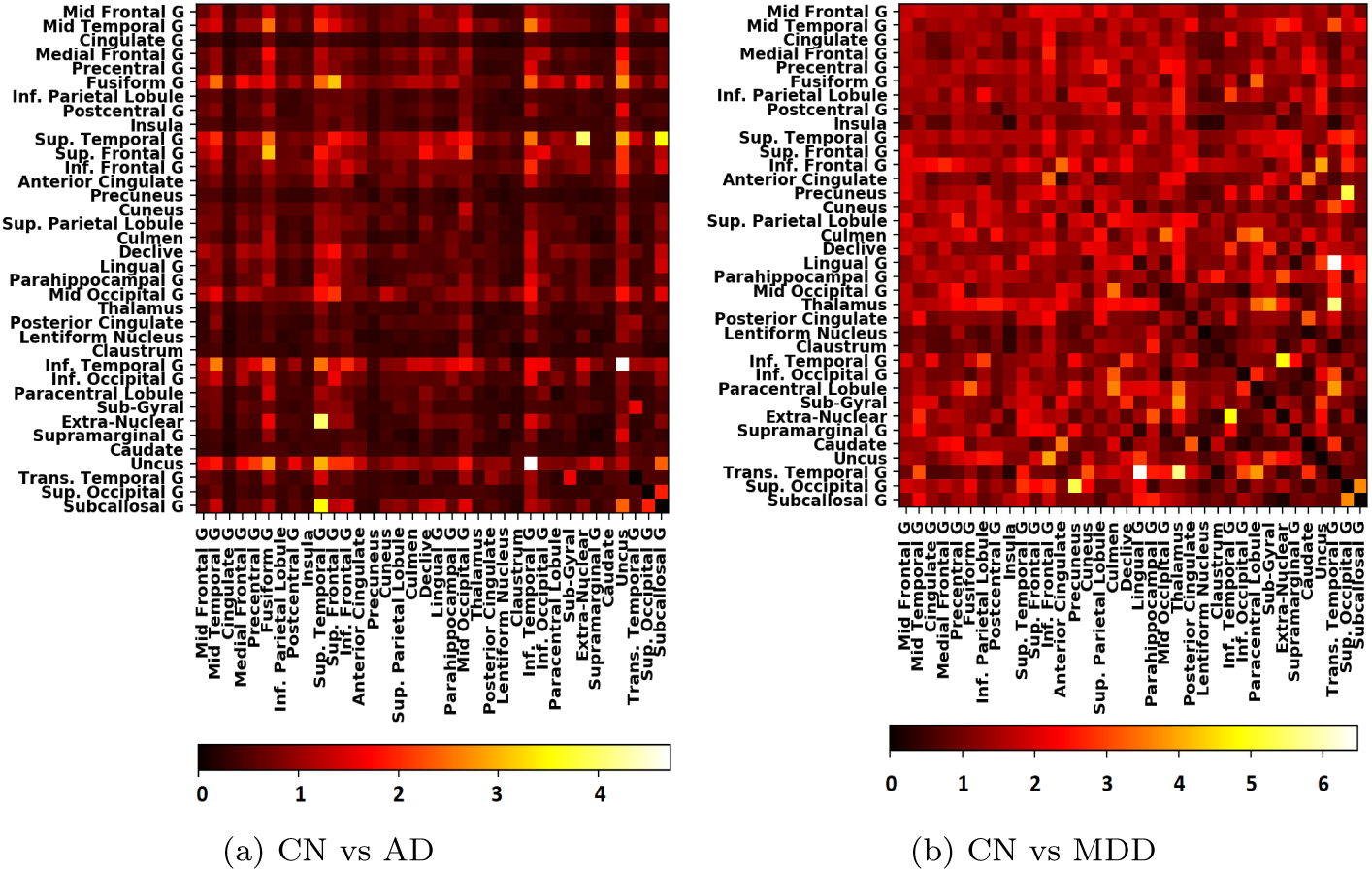
The salience scores of functional connections between brain regions derived while classifying normal and diseased participants for (a) Alzheimer’s Disease; and (b) Major Depressive Disorder. Mid: Middle, Inf: Inferior, Sup: Superior, G: Gyrus

For all the diseases, ROI in the inferior temporal gyrus and paracentral lobule were common among the salient regions. The other most distinctive ROI for MDD were located in the subcallosal gyrus, supplementary motor area, the medial frontal gyrus, the parahippocampal gyrus and the posterior cingulate (Figure 5b); for ADHD in the postcentral gyrus, inferior frontal gyrus, transverse temporal gyrus, superior occipital gyrus, the insula and the medial frontal gyrus (supplementary figure S3(a)). For both AD and MCI, the salient regions were found to be in the cerebellum, temporal pole middle, frontal superior medial, parahippocampal and fusiform gyrus (Figure 5a and supplementary figure S3(b), respectively). For ASD, salient regions were found in the inferior temporal lobe, the rectus, heschl’s gyrus, cerebellum, and the paracentral lobule (supplementary figure S3(c)).

**Figure 5:**
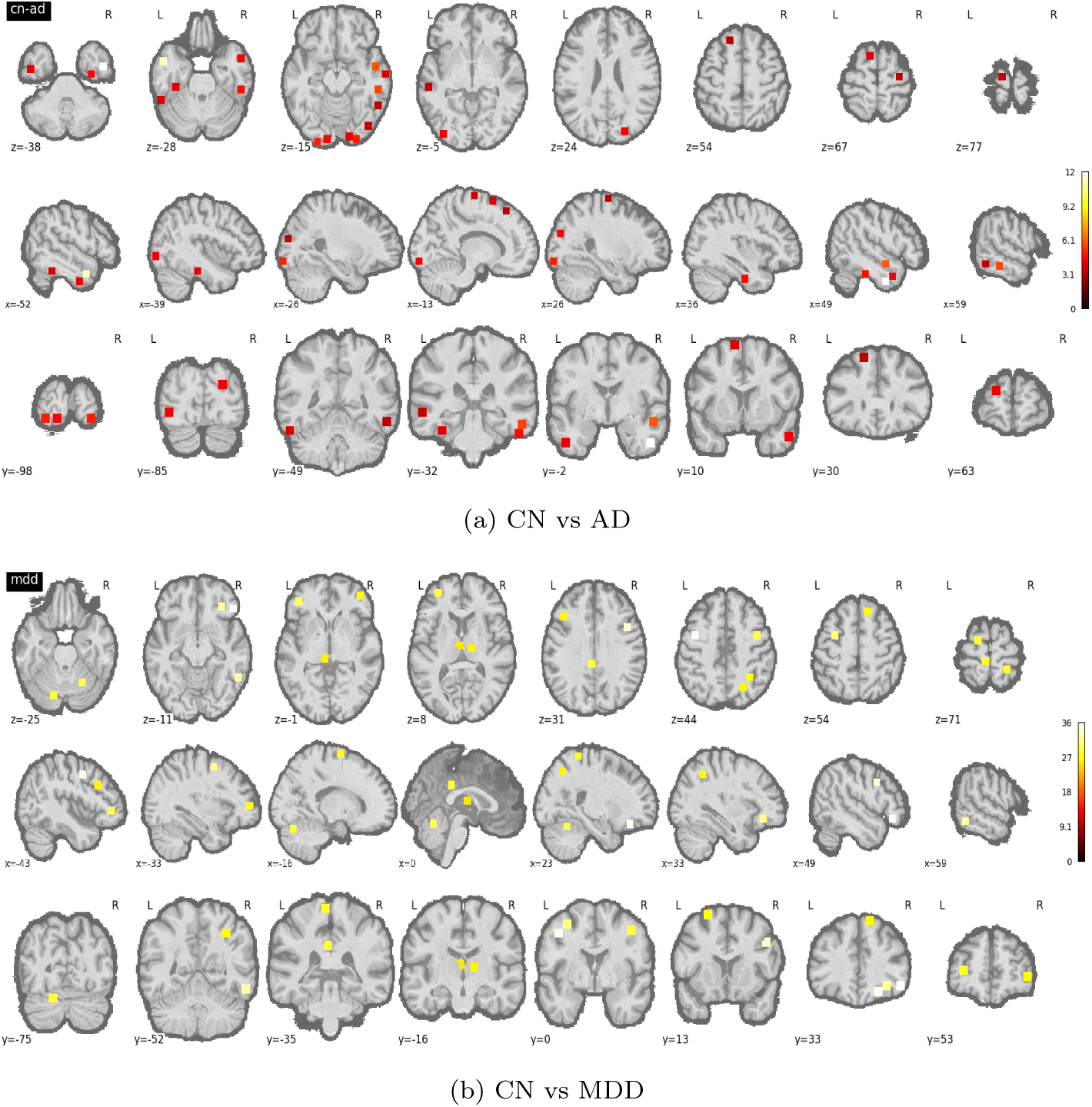
The axial, sagittal and coronal views of top 10% salient ROI differentiating normal and diseased patients for (a) Alzheimer’s Disease, and (b) Major Depressive Disorder.

## 4. Discussion

### 4.1. DNN models for full feature set

As seen in table 1, the feedforward DNN using the entire feature set is generally able to achieve better classification accuracies than CNNs and SVMs. The rather high standard deviation is attributable to the size of the datasets [57], which typically do not contain more than a few hundred scans. The low accuracy for ADHD is attributable to the mismatch between the age of the subjects and the age of the subjects used to derive the Power atlas [35]. Also, more subtle connectivity differences with respect to CN subjects makes it harder to classify MCI as compared to AD.

We found that CNNs and DNNs perform consistently better than SVMs. However, the difference in performance between CNNs and feedforward DNNs are subtle: except for CN-AD, feedforward DNNs did better than CNNs but the difference is rarely more than 1%. From these results, CNNs - even the customised ones - do not seem to be getting any additional significant information for functional connectivity. Although CNNs have desirable features such as parameter sharing, they only capture information within their local receptive field (usually a small square-shaped subset, or a row or column along a matrix). Such information is not sufficient to capture the functional connectivity present in brains as these receptive fields do not consider global connectivity patterns. On the other hand, the fact that CNNs have fewer parameters than feedforward DNNs shows the possibility of redundancies in feedforward DNNs which makes it important to eliminate accessory nodes from feedforward DNNs. With LEAN, we are able to get a leaner model while methodically removing less salient features and nodes (instead of being constricted by the spatial limitations of local receptive fields).

### 4.2. LEAN + CLIP: high accuracy with few parameters

From the results in table 2, we observe that LEAN has the largest accuracy drop but the smallest number of remaining parameters; but, ‘CLIP + LEAN (Inputs Only)’ results in the small drop in accuracy but retains the largest number of parameters. However, ‘CLIP + LEAN’ gives the best of both worlds - the number of weights retained is close to levels from LEAN, but the accuracy drop is similar to that of ‘CLIP + LEAN (Inputs Only)’. An implication of our results is the presence of correlated features in functional connectomes, which can be exploited to reduce input feature set in functional/structural connectomes. In our case, we observed that LEAN led to drastic reduction in the input feature set, thereby removing sets of correlated features entirely. Our results show that adding a subset of the correlated features in ‘CLIP + LEAN’ (and also in ‘CLIP + LEAN (Inputs Only)’) led to an improvement in the performance of the classifier.

Crucially, our results show that there is a large amount of redundancy in neural network models. Our experiments with quantification of overfitting and time taken per epoch for the proposed model show that the proposed model not only leads to a reduction in overfitting but is also faster. The latter was expected since there is a significant reduction in the number of trainable network parameters. Feedforward networks have been used by researchers in the past [19, 25, 27] to perform classification on functional connectivity data. However, these works often use the full set of features and overfitting is a key limitation in such approaches. In our work, we have proposed a way to reduce the effects of overfitting and showed how even as it leads to a drastic reduction of parameters, the drop in accuracy is minimal. Thus, just a simple feedforward DNN is sufficient to capture crucial patterns in the data to classify healthy and diseased brains. Adding more nodes in the hidden layers does not improve model accuracy much (and in some cases such as CN-AD classification, doing so even leads to poorer generalization).

Although we have only tested our approaches on neuroimaging datasets, they are also widely applicable to other datasets. LEAN (Algorithm 1) is applicable in settings where the number of features greatly outnumber the size of the dataset, or in cases where the dataset is small and will benefit from using fewer features so as to increase generalizability of the model. CLIP (Algorithm 2) is applicable to any datasets where there are features that have strong correlations with each other.

### 4.3. Identified salient features are clinically relevant

For our algorithm to perform well, it is important that the features classified as salient by our decoder have clinical relevance to the respective condition. To verify this, we compare the results of our decoder with results from previous studies finding disease biomarkers.

For AD/MCI, previous studies have consistently pointed out to changes in the hippocampal and medial temporal lobe [9, 11, 52]. Accordingly, we found alterations in the inferior temporal, parahippocampal [58], fusiform [9], and paracentral lobule [52]. We observed that the most salient connections were from the uncus for both AD and MCI, an anterior extremity of the parahippocampal gyrus known for disruption during both AD [59] and MCI [65]. Multiple functional networks such as the fronto-parietal task-control network, involved in attention and emotion regulation; the default mode network, for internal medication; the dorsal attention network, for directing external attention; and the salience network, which helps in emotion processing or monitoring salient events have been implicated in MDD [23, 46]. Deficits in cognitive control can be traced to the anomalies in the fronto-parietal task-control network, whereas too much internal rumination (and lesser engagement with the external world) may be reflected in the aberrant connectivity of the default mode network with other goal-directed networks. The amygdala [26, 36] (part of the subcallosal region), regions in the inferior temporal [56], the medial frontal lobe [56], posterior cingulate cortex [63] and supplementary motor area [64] have been implicated in MDD. In [61], disruption between the connections between regions in the thalamus and the transverse temporal gyrus were reported, which is consistent with our results.

Multiple studies have reported anomalies in the default-mode and dorsal attention networks in children and adults suffering with ADHD suggesting altered functional connectivity with attention and cognitive processing [45, 51, 53, 55]. The postcentral gyrus and paracentral lobule areas involved in motor functioning [45, 51], medial and inferior frontal gyrus [39, 45], superior occipital gyrus [55], and the insula [53] have been found to be different for children/young adoloscents having ADHD, which are consistent with our results.

Likewise, the frontal orbital regions and Heschl’s gyrus are involved in sensory integration, speech processing and decision-making [24, 37], the putamen responsible for focusing attention [41], the cerebellar (and vermis) involved in motor regulation [3] have been previously identified as biomarkers for ASD.

## 5. Conclusion

In summary, we have proposed two algorithms - LEAN and CLIP - to reduce overfitting in DNNs making them more generalizable than models that use the entire feature set. Our approach leads us to an optimal neural network architecture in a single shot that is more efficient than previous methods that relied on recursive removal of features and nodes. Furthermore, via CLIP, the approach is customised to reduce the effects of correlated features that are present in neuroimaging datasets. Our experiments show that using both ‘LEAN + CLIP’ took into account both redundancy and correlation in input features and gave the best balance between drop in accuracy and reduction in trainable weights. We successfully applied the proposed algorithms on 4 datasets (and 5 different neurological disorders), showing its application in brain decoding and biomarker identification. The proposed approach has applications to both structural and functional neuroimaging datasets and to investigate brain disease.

## Acknowledgement

This work was partially supported by AcRF Tier 1 grant RG 149/17 of Ministry of Education, Singapore. Data collection and sharing for this project was funded by the Alzheimer’s Disease Neuroimaging Initiative (ADNI) (National Institutes of Health Grant U01 AG024904) and DOD ADNI (Department of Defense award number W81XWH-12-2-0012), the International Neuroimaging Datasharing Initiative (INDI) and the Creativity and Affective Neuroscience Lab in the Brain Imaging Center of Southwest University. We would like to thank Dr. Jiang Qiu and Dr. Dongtao Wei from the Creativity and Affective Neuroscience Lab, Brain Imaging Center of Southwest University for providing us the dataset for MDD.

