## Supplementary for "Obtaining leaner deep neural networks for decoding brain functional connectome in a single shot"

Sukrit Gupta*, Yi Hao Chan*, Jagath C. Rajapakse, and the ADNI

May 27, 2020

#### 1 Datasets

##### 1.1 ABIDE

We used neuroimaging datasets from the Autism Brain Imaging Data Exchange (ABIDE) [1]. ABIDE involves 19 sites with over 1000 fMRI scans datasets from ASD and 593 normal controls (age range: 5-64 years) containing phenotypic characterization, particularly in regard to measures of core ASD and associated symptoms. In this study, we used rs-fMRI data of 387 ASD and 436 control subjects. from the Autism Brain Imaging Data Exchange (ABIDE) [1]. We removed subjects with mean framewise displacement greater than 0.2mm and used the data preprocessed using the C-PAC pipeline from the Preprocessed Connectomes Project.

##### 1.2 ADNI

Functional and structural MRI data used from ADNI were obtained from the ADNI database (`adni.loni.usc.edu`). The rs-fMRI data for each subject consisted of 140 or 200 functional volumes, acquired with the following parameters: repetition time (TR) = 3000 ms; echo time (TE) = 30 ms; flip angle = 80°; slice thickness = 3.313 mm; and 48 slices. Results included in this manuscript come from pre-processing performed using fMRIPprep [2]. As there are too few scans available for MCI, we combined early MCI, MCI and late MCI into one category (refer table S2). Subjects classified as AD fulfilled the criteria for AD laid down by National Institute of Neurological and Communicative Disorders and Stroke and the Alzheimer's Disease and Related Disorders Association. Subjects classified as MCI had a memory complaint, objective memory loss measured by education adjusted scores on Wechsler Memory Scale Logical Memory II, absence of dementia and significant levels of impairment in other cognitive domains. The CN subjects did not suffer from depression, cognitive impairment, or dementia. Table S2 gives a summary of the gender and age distribution of the dataset, and also includes metrics to measure cognitive function.

##### 1.3 ADHD

We obtained the preprocessed ADHD data from the consortium of the International Neuroimaging Datasharing Initiative (INDI), which contains aggregated data from collaboration of 8 international imaging sites [3]. We used the data of subjects preprocessed using the Athena pipeline from the New York University Child Study Center and retained the subjects that were classified as normal and ADHD Combined and had no secondary diagnosis.

*These authors contributed equally.

Table S1: ASD subject details, FIQ Score

| | Cognitively Normal (CN) | Austism Spectrum Disorder (ASD) |
|---|---|---|
| No. of subjects | 436 | 387 |
| Male/Female | 354/82 | 341/46 |
| Age | 16.40 ± 6.97 | 17.86 ± 9.11 |
| FIQ Score | 111.25 ± 12.51 | 105.70 ± 16.76 |

Table S2: ADNI subject details, including Mini-Mental State Examination (MMSE) and Clinical Dementia Rating (CDR) scores

| | Cognitively Normal (CN) | Mild Cognitive Impairment (MCI) | Alzheimer's Disease (AD) |
|---|---|---|---|
| No. of subjects/scans | 49/196 | 90/356 | 29/103 |
| Male/Female | 22/27 | 46/44 | 15/14 |
| Age | 76.6 ± 5.5 | 74.4 ± 6.7 | 75.4 ± 8.2 |
| MMSE score | 28.9 ± 1.1 | 27.3 ± 2.4 | 21.2 ± 2.6 |
| CDR score | 0.13 ± 0.3 | 0.68 ± 0.4 | 1.13 ± 0.6 |

Table S3: ADHD subject details, Conners' Parent Rating Scale-Revised, Long version (CPRS-LV)

| | Cognitively Normal (CN) | Attention Deficit Hyper. Disorder (ADHD) |
|---|---|---|
| No. of subjects/scans | 99/184 | 123/212 |
| Male/Female | 48/51 | 97/26 |
| Age | 12.15 ± 3.13 | 11.13 ± 2.69 |
| CPRS-LV | 45.2 ± 6.01 | 71.1 ± 8.71 |

##### 1.4 MDD

We used the preprocessed data provided by the Creativity and Affective Neuroscience Lab (led by Dr. Jiang Qiu), Brain Imaging Center of Southwest University. The dataset contained rs-fMRI scans collected from 3T MRI scanners for an 8-min period for 289 diseased and 168 CN subjects at Xinan (First Affiliated Hospital of Chongqing Medical School in Chongqing, China). The participants were diagnosed according to the Diagnostic and Statistical Manual of Mental Disorder-IV criteria for MDD and depression severity and symptomatology were evaluated using the Hamilton Depression Rating Scale (HAMD, 17 items) [4] and the Beck Depression Inventory (BDI) [5].

#### 2 Functional MRI Preprocessing

##### 2.1 ABIDE

In this study, we used rs-fMRI data of 387 ASD and 436 control subjects from ABIDE [1], a consortium of brain imaging data for sharing within the scientific community. We removed subjects with mean framewise displacement greater than 0.2mm and used the data preprocessed using the C-PAC pipeline from the Preprocessed Connectomes Project. The preprocessing of the fMRI data includes correction of slice timing, realignment of motion, voxel intensity normalization, nuisance signal removal and band-pass filtering.

##### 2.2 ADNI dataset

For each of the fMRI scan for a subject, the following preprocessing was performed. First, a reference volume and its skull-stripped version were generated and the BOLD reference was co-registered to the T1w reference using `bbregister` (FreeSurfer). Head-motion parameters with respect to the BOLD reference (transformation matrices, and six corresponding rotation and translation parameters) are estimated before any spatiotemporal filtering using `mcflirt` [6]. BOLD runs were slice-time corrected using `3dTshift` from AFNI [7]. The BOLD time-series were resampled onto their original, native space by applying a single, composite transform to correct for head-motion and

Table S4: MDD subject details, Hamilton Depression Rating Scale (HAM-D)

| | Cognitively Normal (CN) | Major Depressive Disorder (MDD) |
|---|---|---|
| No. of subjects | 168 | 289 |
| Male/Female | 66/102 | 99/190 |
| Age | 31.01 ± 11.79 | 38.74 ± 13.65 |
| HAM-D score | - | 20.78 ± 5.87 |

susceptibility distortions.The BOLD time-series were resampled to `MNI152NLin2009cAsym` standard space, generating a preprocessed BOLD run in MNI152NLin2009cAsym space. Principal components are estimated after high-pass filtering the preprocessed BOLD time-series. The head-motion estimates calculated in the correction step were also placed within the corresponding confounds file. The BOLD time-series, were resampled to *fsaverage5* surfaces with *a single interpolation step* by composing all the pertinent transformations (i.e. head-motion transform matrices, susceptibility distortion correction when available, and co-registrations to anatomical and template spaces). Gridded (volumetric) resamplings were performed using antsApplyTransforms (ANTs), configured with Lanczos interpolation to minimize the smoothing effects of other kernels [8]. Non-gridded (surface) resamplings were performed using `mri_vol2surf` (FreeSurfer).

##### 2.3 ADHD dataset

Athena's rs-fMRI pipeline involved removing the first four volumes, slice timing correction, realignment for motion correction and linear transformation between the subject's functional and structural MRI image. This was followed by computation of WM and CSF signals and regressing 6 head motion parameters and variation due to physiological noise, head motion, and scanner drifts from the time series. Band-pass filtering and spatial smoothening with a 6 mm FWHM Gaussian filter was applied further. Further details can be seen in [3].

##### 2.4 MDD

We used the preprocessed data provided by the Creativity and Affective Neuroscience Lab (led by Dr. Jiang Qiu), Brain Imaging Center of Southwest University. The preprocessing was performed using DPARSF [9] (http://restfMRI.net), which is a toolbox based on the SPM8 software package. They discarded the first 10 EPI scans and the remaining scans underwent slice timing correction, motion correction, spatial normalization to standard Montreal Neurological Institute (MNI) space followed by spatial smoothing. To remove spurious correlations band-pass temporal filtering (0.01–0.1 Hz), nuisance signal removal from the ventricles and deep white matter and regression of 24 head motion parameters was applied. They also performed scrubbing movement correction Power et al. [10] and removed subjects with mean framewise displacement greater than 10%. For more details on preprocessing, please refer to [11].

#### 3 Finding optimal hyperparameters for the models

For the FFN, we varied the number of hidden layers and the number of neurons in each hidden layer. For the CNN we tried different architectures (with different filters), and varied the number of filters in each layer. The number of filters (in case of CNN) and the number of neurons (in case of FFN) were fixed such that the number of trainable parameters were approximately the same. For FFN we varied the number of hidden layers from 1 to 3, and the number of hidden layer neurons from 5 to 1000. For SVM models, we varied the kernel type, the $C$ and the parameter $\gamma$.

For CNN, we defined 3 types of convolution layers:

1. A 'closeness layer' containing convolution filters of size $2 \times 2$.
2. A layer containing filters of shape $1 \times |ROI|$.
3. A layer containing filters of shape $|ROI| \times 1$.

We tried 2 different architectures, where we tried all the layers listed above (which we termed as funcNetCNN), and removed the closeness layer (which we termed as funcNetCNN_C). This was based on the previously proposed architectures by [12, 13]. The number of filters in the layers were varied such that they had approximately the same number of parameters as the other CNN models and the corresponding FFN models. The convolution layers were followed by a fully connected layer of 32 neurons and a softmax layer for classification. The accuracies for different models (FFN, SVM and the CNN) are given in Figure S1.

#### 4 Selected features for ADHD, MCI and ASD by LEAN and CLIP

We analyse the importance of the functional connections by using the salience scores and the importance of ROI by computing the sum of salience score of the functional connections incident on each of the ROI. We map the ROI of the Power atlas [14] to the regions in the Crossley atlas [15] on the basis of Euclidean distance and assign anatomical labels to Power atlas ROI. We plot the salience score of the connections in Figure S2 and the anatomical locations of the top 10% salient ROI are visualised using Nilearn S3.

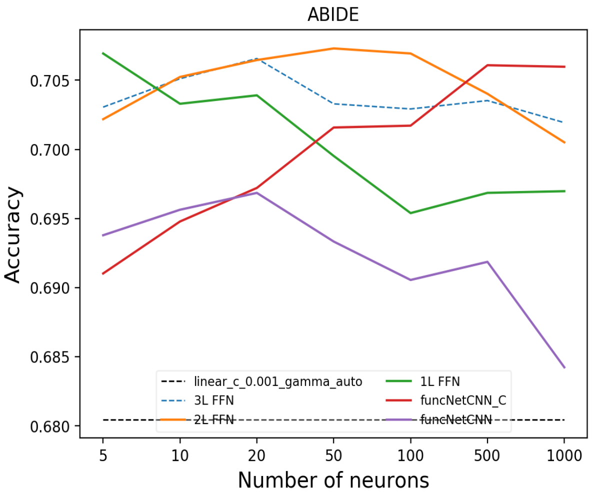


(a) CN vs ASD

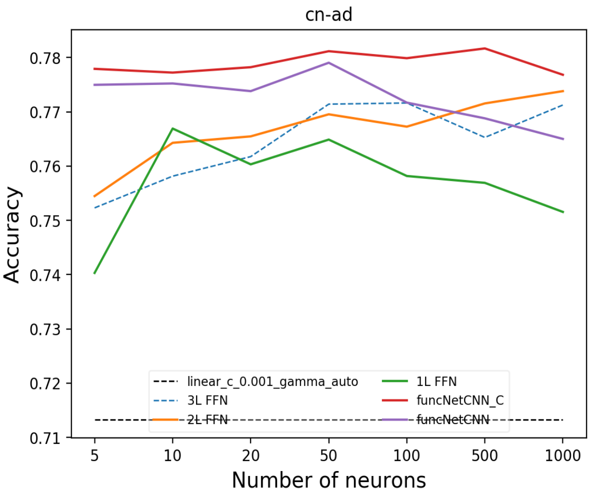


(b) CN vs AD

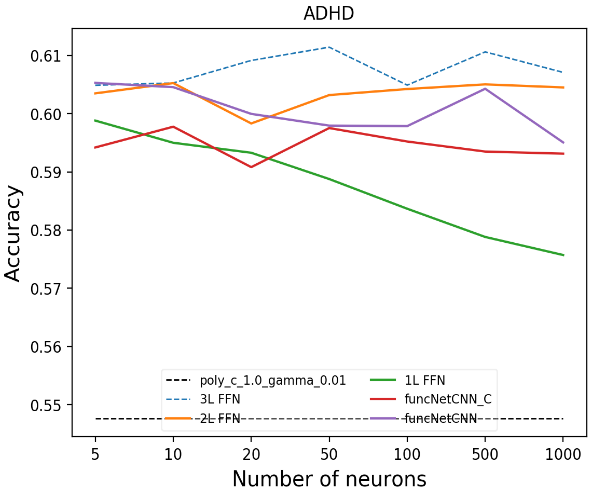


(c) CN vs ADHD

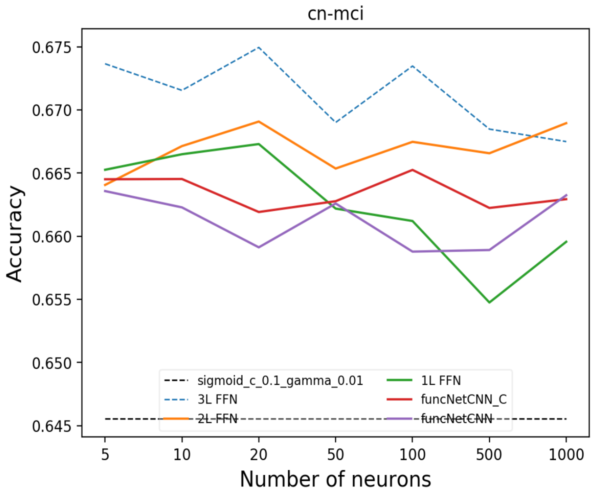


(d) CN vs MCI

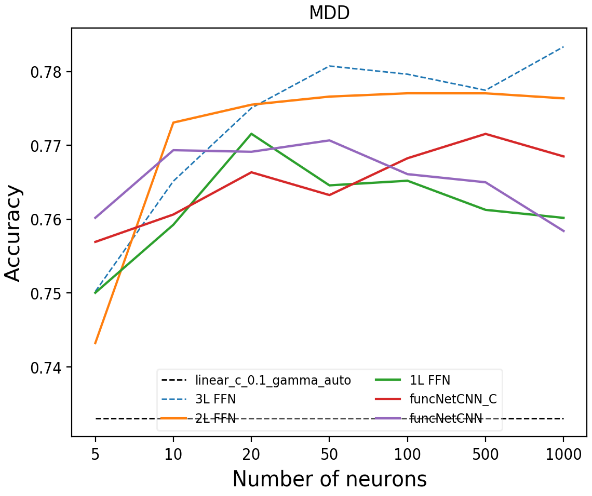


(e) CN vs MDD

Figure S1: Accuracy plots of SVM, FFN and CNN models applied on the various neuropsychiatric diseases. The black dotted lines show the results for SVM (a straight line since there are no neurons in SVMs) and the legend shows the exact configuration used).

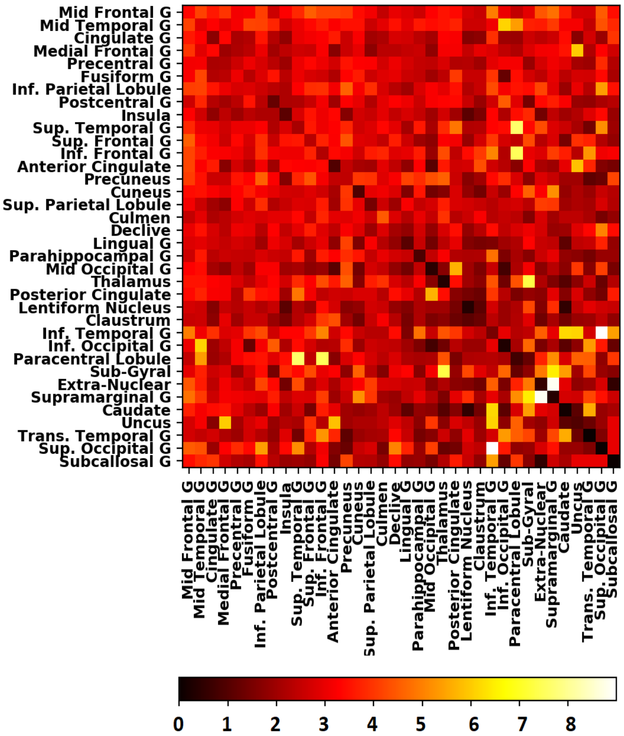


(a) CN vs ADHD

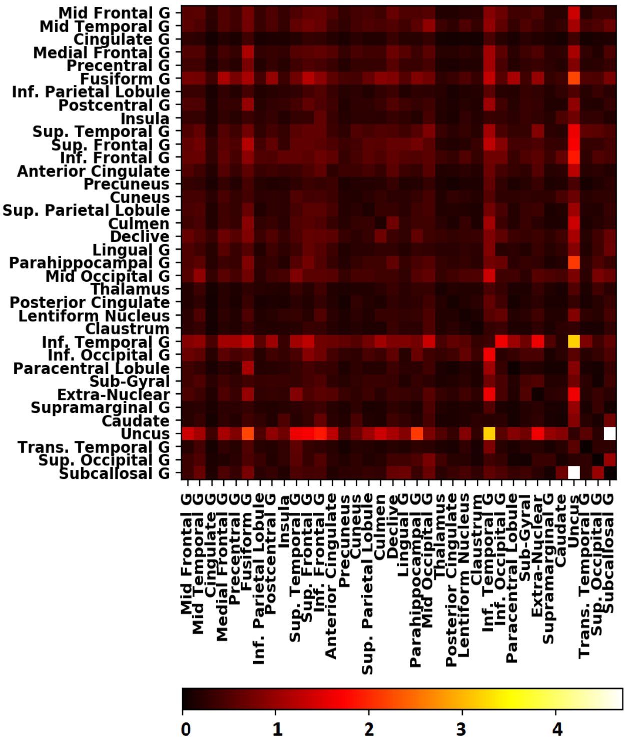


(b) CN vs MCI

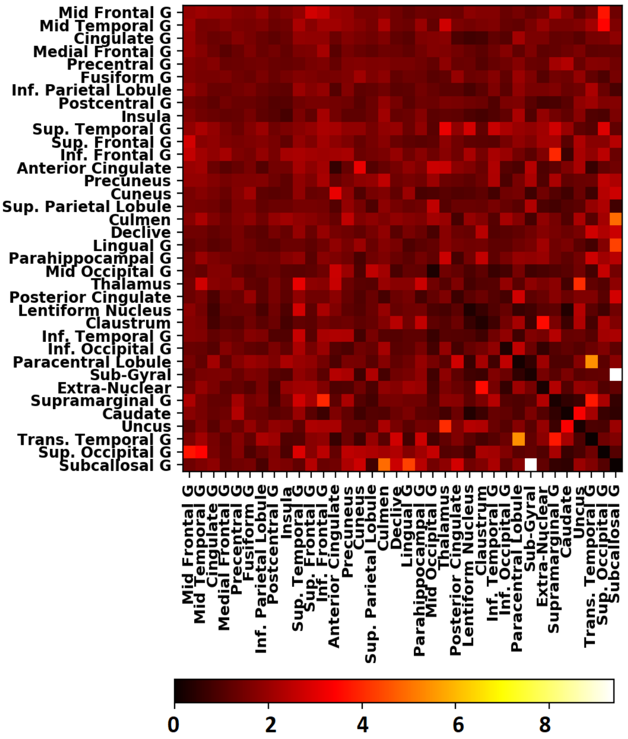


(c) CN vs ASD

Figure S2: The salience scores of functional connections between brain regions derived while classifying normal and diseased participants for (a) attention deficit hyperactivity disorder; and (b) mild cognitive impairment. Mid: Middle, Inf: Inferior, Sup: Superior, G: Gyrus

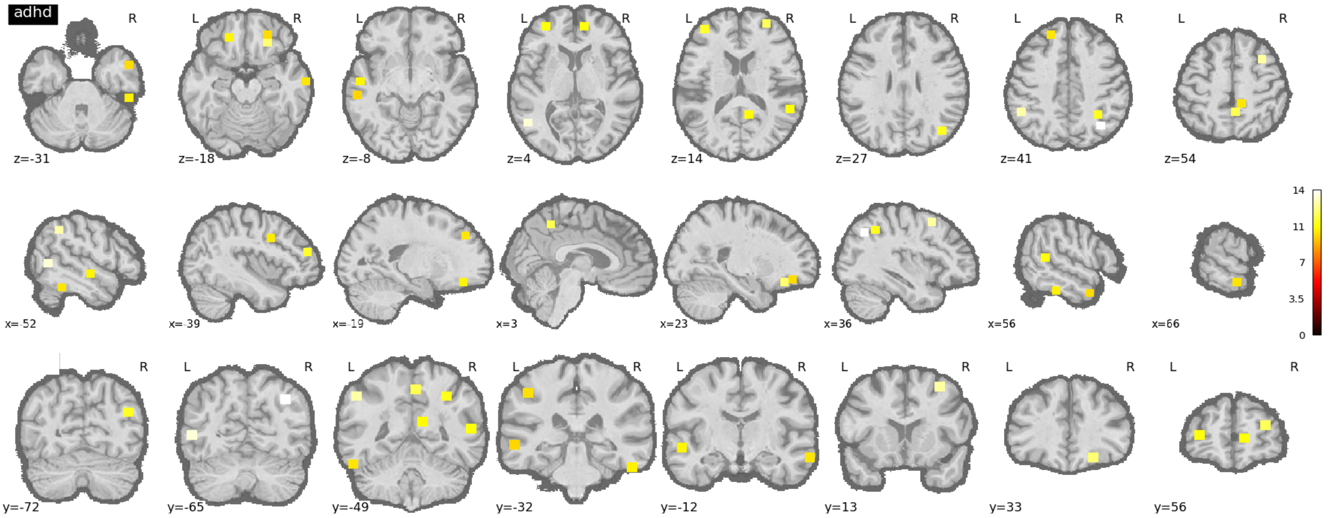


(a) CN vs ADHD

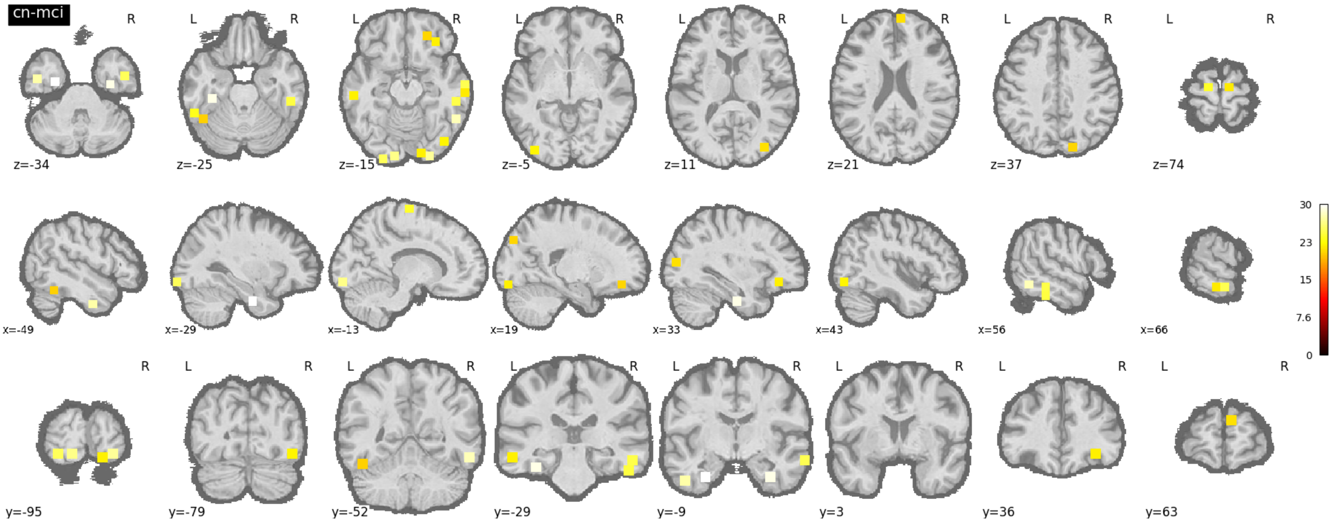


(b) CN vs MCI

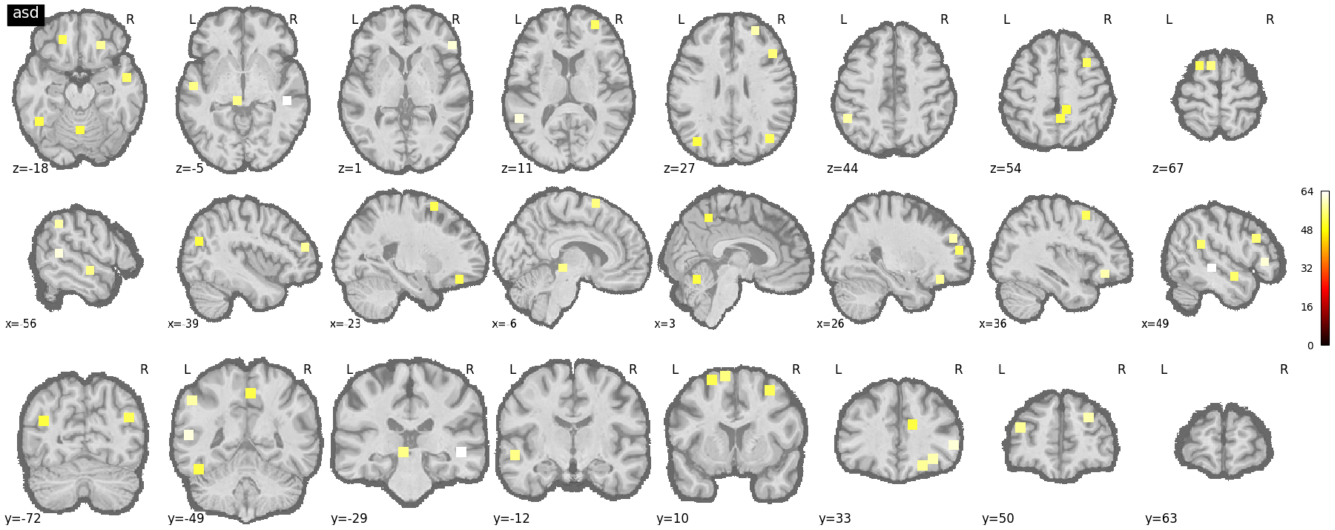


(c) CN vs ASD

Figure S3: The axial, sagittal and coronal views of top 10% salient ROI differentiating normal and diseased patients for (a) attention deficit hyperactivity disorder, (b) mild cognitive impairment, and (c) autism spectrum disorder.

Table S5: Best architectures found of SVM Models on the neuroimaging datasets

| Dataset | SVM Model (Kernel, C, $\gamma$) | CNN Model | FFN Model (hidden layer config) |
|---|---|---|---|
| ABIDE | Linear , 0.001, - | funcNetCNN_C1000 | 50-10 |
| AD | Linear, 0.001, - | funcNetCNN_C50 | 50-32-32 |
| ADHD | Polynomial, 1.0, 0.01 | funcNetCNN_500 | 50-32-32 |
| MCI | Sigmoid, 0.1, 0.01 | funcNetCNN_C100 | 20-32-32 |
| MDD | Linear, 0.1, - | funcNetCNN_50 | 1000-64-32 |

### 5 Acknowledgment

The ADNI was launched in 2003 as a public-private partnership, led by Principal Investigator Michael W. Weiner, MD. The primary goal of ADNI has been to test whether serial magnetic resonance imaging (MRI), positron emission tomography (PET), other biological markers, and clinical and neuropsychological assessment can be combined to measure the progression of mild cognitive impairment (MCI) and early Alzheimer's disease (AD).

ADNI is funded by the National Institute on Aging, the National Institute of Biomedical Imaging and Bioengineering, and through generous contributions from the following: AbbVie, Alzheimer's Association; Alzheimer's Drug Discovery Foundation; Araclon Biotech; BioClinica, Inc.; Biogen; Bristol-Myers Squibb Company; CereSpir, Inc.; Cogstate; Eisai Inc.; Elan Pharmaceuticals, Inc.; Eli Lilly and Company; EuroImmun; F. Hoffmann-La Roche Ltd and its affiliated company Genentech, Inc.; Fujirebio; GE Healthcare; IXICO Ltd.; Janssen Alzheimer Immunotherapy Research & Development, LLC.; Johnson & Johnson Pharmaceutical Research & Development LLC.; Lumosity; Lundbeck; Merck & Co., Inc.; Meso Scale Diagnostics, LLC.; NeuroRx Research; Neurotrack Technologies; Novartis Pharmaceuticals Corporation; Pfizer Inc.; Piramal Imaging; Servier; Takeda Pharmaceutical Company; and Transition Therapeutics. The Canadian Institutes of Health Research is providing funds to support ADNI clinical sites in Canada. Private sector contributions are facilitated by the Foundation for the National Institutes of Health (www.fnih.org). The grantee organization is the Northern California Institute for Research and Education, and the study is coordinated by the Alzheimer's Therapeutic Research Institute at the University of Southern California. ADNI data are disseminated by the Laboratory for Neuro Imaging at the University of Southern California.
